# Backward alpha oscillations shape perceptual bias under probabilistic cues

**DOI:** 10.1101/2025.01.14.632925

**Authors:** Luca Tarasi, Andrea Alamia, Vincenzo Romei

## Abstract

Predictive coding theory suggests that prior knowledge is crucial for optimizing human decision-making, with recent studies emphasizing the role of alpha-band oscillations in this process. Here, we employed a traveling waves approach to investigate how alpha oscillations integrate prior expectations during a perceptual decision-making task. Our findings demonstrated that expectation-based knowledge triggers the propagation of alpha traveling waves from frontal to occipital areas, with this increase associated with enhanced modulation of brain regions involved in stimulus processing and directly linked to prior-driven bias at the behavioral level. Moreover, participants who relied more on prior expectations exhibited stronger top-down signaling, whereas those who focused on sensory input showed a contrasting forward signaling pattern. These results highlight the role of alpha-band traveling waves in predictive mechanisms, suggesting that rhythmic interactions across brain regions facilitate this process and contribute to inter-individual differences in its implementation.

## Introduction

The brain’s capacity to generate predictions plays a pivotal role in perception and decision-making ^1^. Predictive coding theory proposes that the brain continuously generates expectations about incoming sensory information based on prior knowledge, forming an “internal model” to interpret the sensory world ^2^. The predictive machinery relies heavily on top-down information flow, whereby higher-order cortical areas send signals to lower-level regions ^3,4^. Such signals modulate sensory systems to refine perceptual experiences based on the probabilistic structure of the environment. This modulation, in turn, leads to shifts in perceptual parameters, including reaction time, decision criteria, and decision confidence ^5–10^. Brain oscillations, particularly in the alpha band (8-14 Hz), are understood to play a critical role in coordinating these top-down modulations. Traditionally associated with attentional and inhibitory functions ^11–18^, alpha activity is hypothesized to also support predictive processing by conveying top-down information flow across the cortical hierarchy ^19^. Specifically, alpha rhythms appear to adapt decisional outcomes in alignment with environmental expectations, modulating responses according to probabilistic cues and fostering perceptual and behavioral adaptations. For example, we demonstrated that the modulation of parieto-occipital alpha amplitude underpins probabilistic cue integration into perceptual processing ^20^. Additionally, Kloosterman et al. ^21^ showed that posterior alpha amplitude modulation serves to strategically bias evidence accumulation during perceptual tasks. Moreover, it accounts for the modulation of both criterion ^22^ and confidence levels in the response ^23–25^.

Crucially, alpha oscillations are not static but can flow as “travelling waves” across cortical areas, propagating from one region to another in a specific direction (e.g., anterior to posterior or vice versa). These travelling waves are thought to facilitate efficient signal transmission between brain regions, supporting broad cortical communication ^26–28^. Within a predictive coding framework, alpha travelling waves may convey prediction-based information from higher-order brain regions toward areas specializing in sensory processing, thereby preparing the perceptual system for upcoming stimuli. The directionality of these waves would be particularly revealing in this domain: while forward-directed waves (anterior-to-posterior) are particularly enacted during sensory stimulation ^11,29,30^, backward-directed waves (posterior-to-anterior) may reflect a predictive signal moving from higher-order areas to sensory area in order to shape responses based on expectations.

Here, we investigated the role of alpha travelling waves in incorporating predictive information into perceptual decision-making. We used a visual detection task in which human observers viewed visual stimuli preceded by probabilistic cues that suggested the likelihood of target appearance. Drawing from our previous research ^20,31^, we hold that these cues significantly influence participants’ decision criteria, serving as a robust means of understanding how probabilistic expectations are integrated into perceptual judgments. Using EEG, we analyzed alpha oscillations as travelling waves to understand how this predictive information affects perception in the task at hand. Specifically, we focused on alpha wave directionality in brain regions contralateral to the expected stimulus. We aimed to determine whether backward-travelling alpha waves could serve as carriers of probabilistic information, thereby modulating participants’ perceptual decisions accordingly. Furthermore, we investigated whether individual differences in the effects of priors on perception were underpinned by distinct modulation of travelling waves.

Our findings reveal a significant increase in backward alpha waves following cue induction in the contralateral hemisphere during the prestimulus period. This contralateral lateralization suggests that backward alpha waves serve as a means of conveying predictive information to the brain regions responsible for stimulus processing, effectively preparing the perceptual system for stimulus anticipation. Additionally, our results demonstrate a robust link between the modulation of decision criteria at the behavioral level and the strength of these backward alpha waves. Participants exhibiting stronger backward-travelling alpha waves showed a greater likelihood of integrating predictive information from the cues into their perceptual decisions, adjusting their decision criteria in alignment with the cue’s predictions.

These results provide compelling evidence for the role of backward alpha travelling waves in integrating probabilistic cues into perceptual decision-making. By supporting the transmission of predictive information to sensory-processing regions, alpha waves appear to facilitate a dynamic interaction with environmental cues. This dynamic modulation supports strategic decision-making, underscoring the crucial function of alpha rhythms in aligning perceptual systems with the brain’s predictive model.

## Results

Eighty participants performed a detection task (Figure 1A). In each trial, a checkerboard pattern appeared in the lower left visual field, which either included isoluminant grey circles within its cells (target trials) or did not (catch trials). Participants were asked to respond using the keyboard to indicate whether they detected the target. Initially, each participant completed an adaptive titration phase similar to the one employed in Tarasi and Romei ^32^ to determine the contrast level of the grey circles that would yield a detection accuracy of 70%. In the subsequent phase, the checkerboards were preceded by a symbolic cue that signalled the likelihood of the target’s appearance. There were three levels of cue probability: a high probability cue indicated a 67% chance of target presence (high probability condition), a low probability cue indicated a 33% chance (low probability condition), and a neutral cue indicated an equal probability (50%) of target presence or absence. The actual target presentation adhered to these probabilities, and participants were explicitly informed that the cues reliably reflected the likelihood of the target’s appearance.

**Figure 1.**
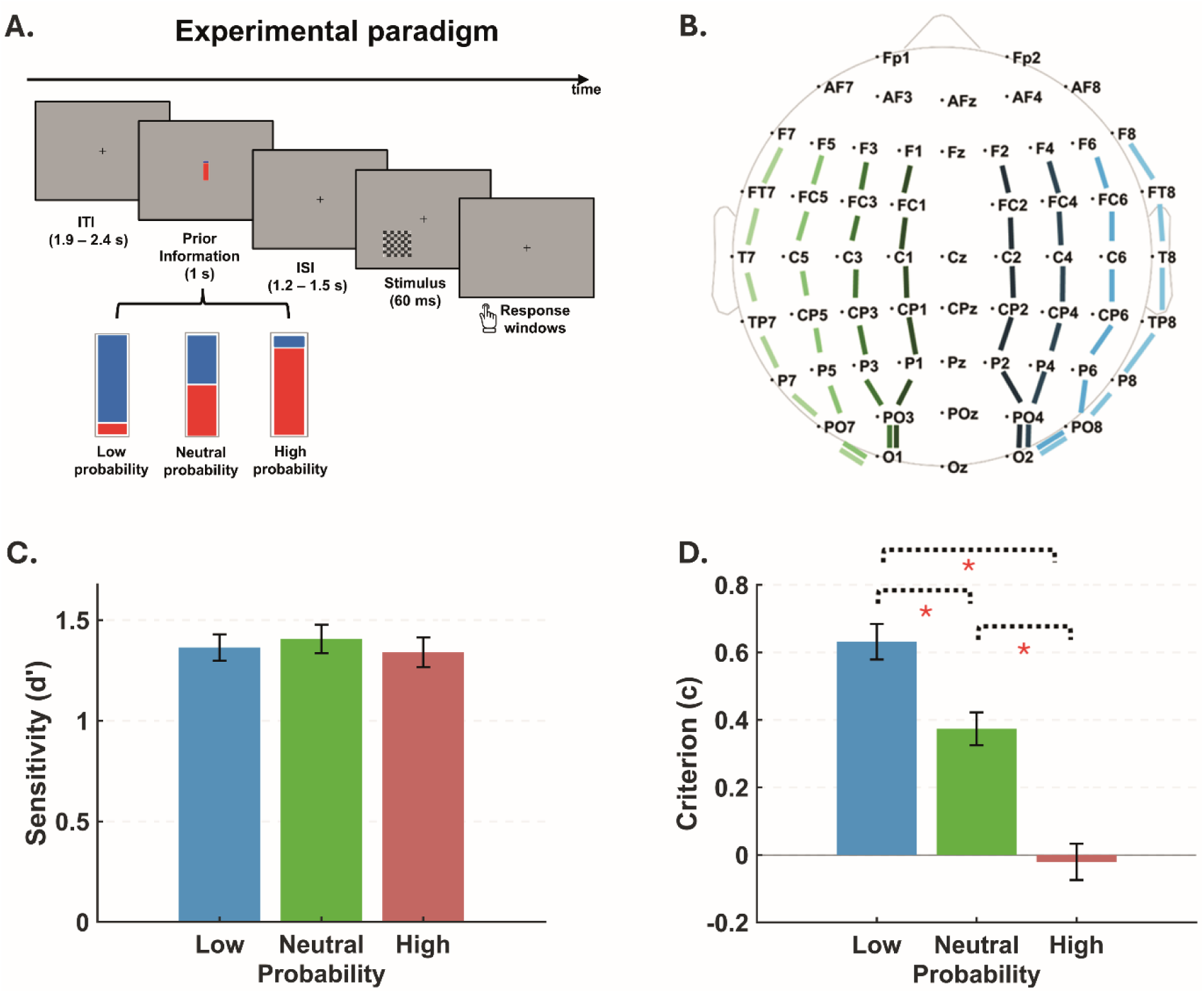
Experimental Setup and Behavioral Results. A) EEG data were collected during a basic visual detection task. Each trial started with a fixation cross, followed by a probabilistic cue in the center of the screen. A checkerboard then briefly appeared (60 ms) in the lower left corner, that could contain grey circles at a set contrast level. The cue was a rectangular bar, with red on the bottom and blue on top, indicating target probability based on the color ratio. The task included three probability levels: high (67% target likelihood), low (33% target likelihood), and neutral (50% chance of target presence or absence). B) Travelling waves were assessed across 8 lines of electrodes positioned along the anterior-posterior axis in the prestimulus period (-600 ms to 0ms). These electrode lines were either contralateral or ipsilateral to the stimulus location. C) Sensitivity and D) Criterion values are shown for trials following low, high, and neutral probability cues. Prior information did not affect sensitivity but had a substantial influence on decision strategies. Specifically, participants used a more liberal criterion after high-probability cues compared to neutral or low-probability cues. Conversely, a more conservative criterion was applied after low-probability cues, relative to neutral cues.

### Probabilistic cue shaped decisional criterion

We computed the signal detection theory indices d’ (sensitivity) and c (criterion) (Green and Swets, 1966) separately for trials preceded by low, high, or neutral probability cues. As demonstrated in the previous publications ^20,31^, the repeated-measures ANOVA revealed no significant effect of the probabilistic cue on *d’* (Figure 1C; F _2,158_ = 1.22; p = 0.30, ηp2 = 0.01). However, the probabilistic cue significantly influenced the criterion (Figure 1D; F_2,158_ = 89.04; p < 0.01, ηp2 = 0.53). Participants adopted a more conservative criterion when the cue indicated a low probability of the target (c _low probability_ = 0.63 ± 0.05), compared to trials with a neutral cue (c _mid probability_ = 0.37 ± 0.05; t_79_ = 7.15, p < 0.01, BF > 1000) or a high probability cue (c _high probability_ = −0.02 ± 0.05; t_79_ = 9.87, p < 0.01, BF > 1000). Moreover, a more liberal criterion was observed when the high probability cue preceded the checkerboard, relative to the neutral cue (t_79_ = −9.78, p < 0.01, BF > 1000).

### Backward travelling waves drive probabilistic cue integration in perceptual decision-making task

We investigated how expectation-like information influences alpha travelling wave patterns. First, we quantified the amount of travelling waves in the contra- and ipsi-lateral hemispheres to the stimulus presentation (Figure 1B). We observed a lateralization effect revealing an increase of contralateral (vs. ipsilateral) backward (but not forward) waves in the alpha-band (Figure 2A).

**Figure 2.**
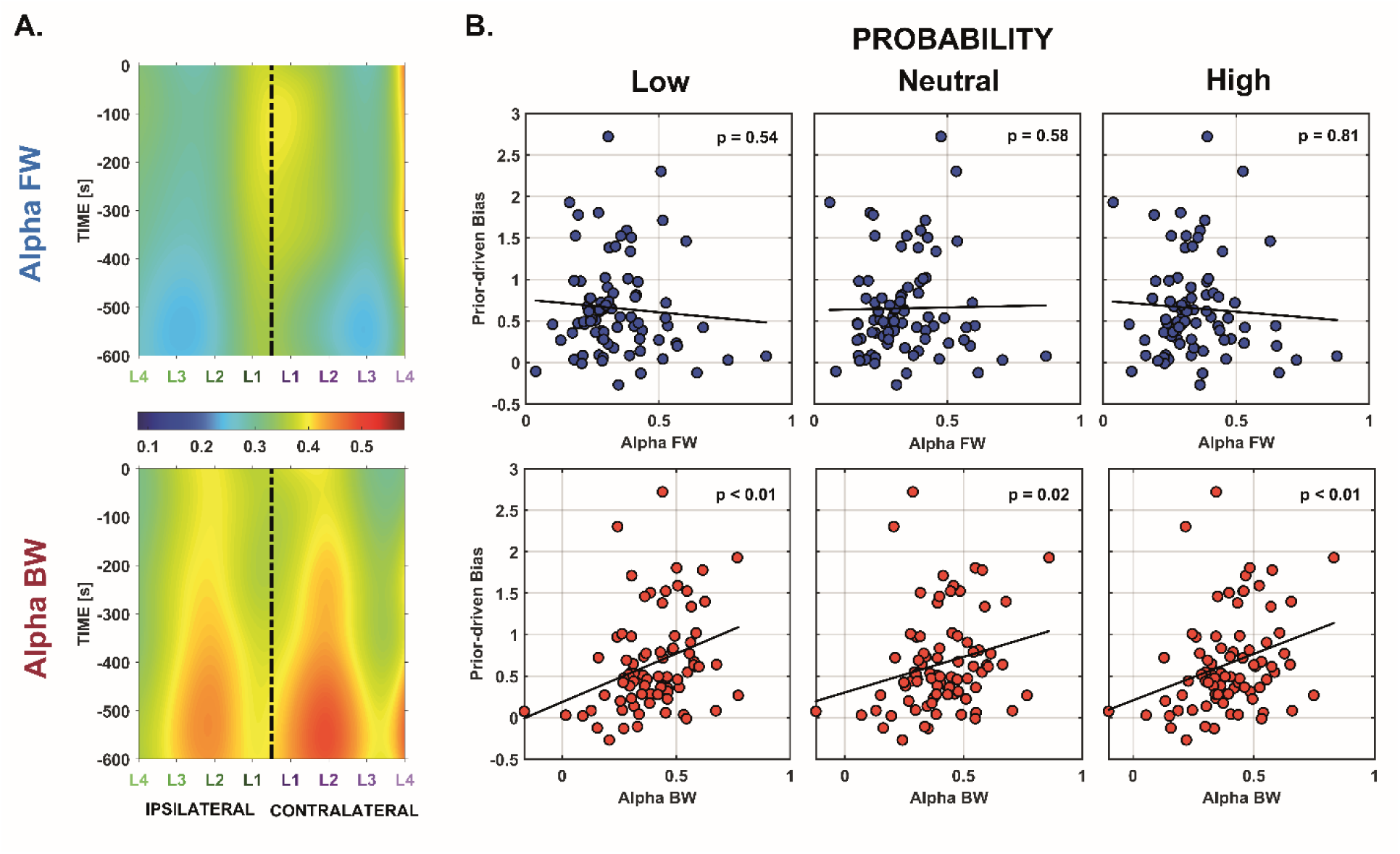
Influence of Expectation-like Information on Travelling Wave Patterns. A) We quantified the dynamics of backward and forward alpha waves in both the contralateral (i.e., right) and ipsilateral (i.e., left) hemispheres relative to stimulus presentation. The accompanying map depicts the magnitude of these waves during the prestimulus period across the specified lines of electrodes, with blue representing low strength and red indicating high strength. A lateralization effect is observable, showing a significant increase in contralateral backward waves in the alpha-band. In contrast, forward waves strength is comparable between the two hemispheres. B) Statistical analysis revealed the presence of an interaction between cue (low,neutral and high probability) and criterion modulation when considering the alpha BW waves in the contralateral hemisphere (F 2,156 = 4.95, p < 0.01, ηp2 = 0.06). Specifically, BW waves in both the low (Spearman = 0.35, p < 0.01) and high (Spearman = 0.32, p < 0.01) probability conditions significantly predicted the magnitude of decisional bias. A similar, though weaker, relationship was also observed in the neutral condition (Spearman = 0.26, p = 0.02). Notably, contralateral alpha forward waves did not predict the degree of criterion modulation in any of the cues considered (all p > 0.54). For visualization purposes, the figure presents the average alpha FW/BW for each subject across contralateral electrodes, as the effect was not specific to any individual electrode line.

We statistically investigated whether the observed alpha traveling waves pattern underpinned the integration of probabilistic cues in the perceptual task. To this end, we conducted two ANCOVA, using CUE (low, neutral, high probability), LINE (1 - 4; distance from the midline) and HEMISPHERE (contralateral, ipsilateral) as factors, with CRITERION SHIFT—an index of cue integration (see methods)—included as a covariate. In the first ANCOVA, BW waves were included as the dependent variable, whereas in the second, FW waves served as the dependent variable.

The first ANCOVA focusing on alpha backward waves (but not when considering theta or beta backward waves, see SI) revealed a significant three-way interaction among CRITERION SHIFT, HEMISPHERE, and CUE (F2,156 = 4.95, p < 0.01, ηp2 = 0.06). To further interpret this result, we examined the relationship between CUE and CRITERION SHIFT separately for each hemisphere. The analysis showed a significant CUE*CRITERION SHIFT interaction in the contralateral hemisphere (F2,156 = 3.71, p = 0.03, ηp2 = 0.05), whereas no significant effect was observed in the ipsilateral hemisphere (F2,156 = 0.14, p = 0.87, ηp2 < 0.01). Post-hoc analyses revealed that the amount of BW alpha waves extracted from the low probability (Pearson = 0.33, p < 0.01; Spearman = 0.35, p < 0.01; skipped Pearson = 0.34, CI = [0.13, 0.53]; skipped Spearman = 0.35, CI = [0.12, 0.56], BF = 9.04) and high-probability conditions (Pearson = 0.30, p < 0.01; Spearman = 0.32, p < 0.01; skipped Pearson = 0.34, CI = [0.14, 0.50]; skipped Spearman = 0.33, CI = [0.10, 0.52], BF = 4.85) significantly predicted the extent of criterion modulation (Figure 2B). The association between BW in the neutral condition and criterion modulation also showed a significant association, although this association was weaker compared to the other conditions (Pearson = 0.23, p = 0.04; Spearman = 0.26, p = 0.02; skipped Pearson = 0.27, CI = [0.07, 0.48]; skipped Spearman = 0.28, CI = [0.06, 0.50], BF = 1.13).

Conversely, in the second ANCOVA focused on FW waves, we found no significant effect of the covariate CRITERION SHIFT nor any interaction with this covariate (all F < 2.67, all p > 0.07, all ηp2 < 0.03).

These findings demonstrate that alpha-band backward traveling waves play a critical role in probabilistic cue integration, with a significant laterality effect driven by top-down activity in the contralateral hemisphere responsible for processing sensory input. This suggests that expectation-like cues modulate neural dynamics, effectively preparing the brain activity of the relevant hemisphere to incoming stimuli in the expected contralateral hemifield.

### Interindividual differences in cue integration strategies are reflected in distinct travelling wave profiles

In our previous research^20^, we identified two distinct groups based on their use of predictive information (see methods and Figure 4A). The amplitude of the posterior alpha oscillations allowed us to intercept these differentiations: participants who exhibited a greater suppression in the amplitude of alpha oscillations in the high-probability vs low-probability condition showed a concurrent strong bias shift (prior-driven individuals), reflecting a more pronounced modulation of their criterion. In contrast, individuals who exhibited a reduced modulation of alpha amplitude showed a dampened criterion shifting (sensory-driven individuals).

Here, we aimed to investigate whether traveling waves patterns could reveal the underlying processes driving individual differences in the integration of expectations during decision-making. To accomplish this, we extracted alpha FW and BW waves (Figure 3A; 3B) from individuals exhibiting above-median cue-driven modulation of their alpha amplitude oscillations (prior-driven group) and compared them to those in a group with lesser modulation (sensory-driven group). We hypothesized that the prior-driven group would exhibit stronger alpha BW activity, whereas the sensory-driven group would display an opposite pattern, characterized by increased alpha FW activity. Importantly, we predicted that these effects would be present in the contralateral hemisphere relative to the stimulus position, compared to the ipsilateral hemisphere. Furthermore, we incorporated the different electrode lines used to extract the traveling waves in the analyses, aiming to investigate whether the hypothesized hemisphere effect was, in turn, modulated by the particular electrode line under examination.

**Figure 3.**
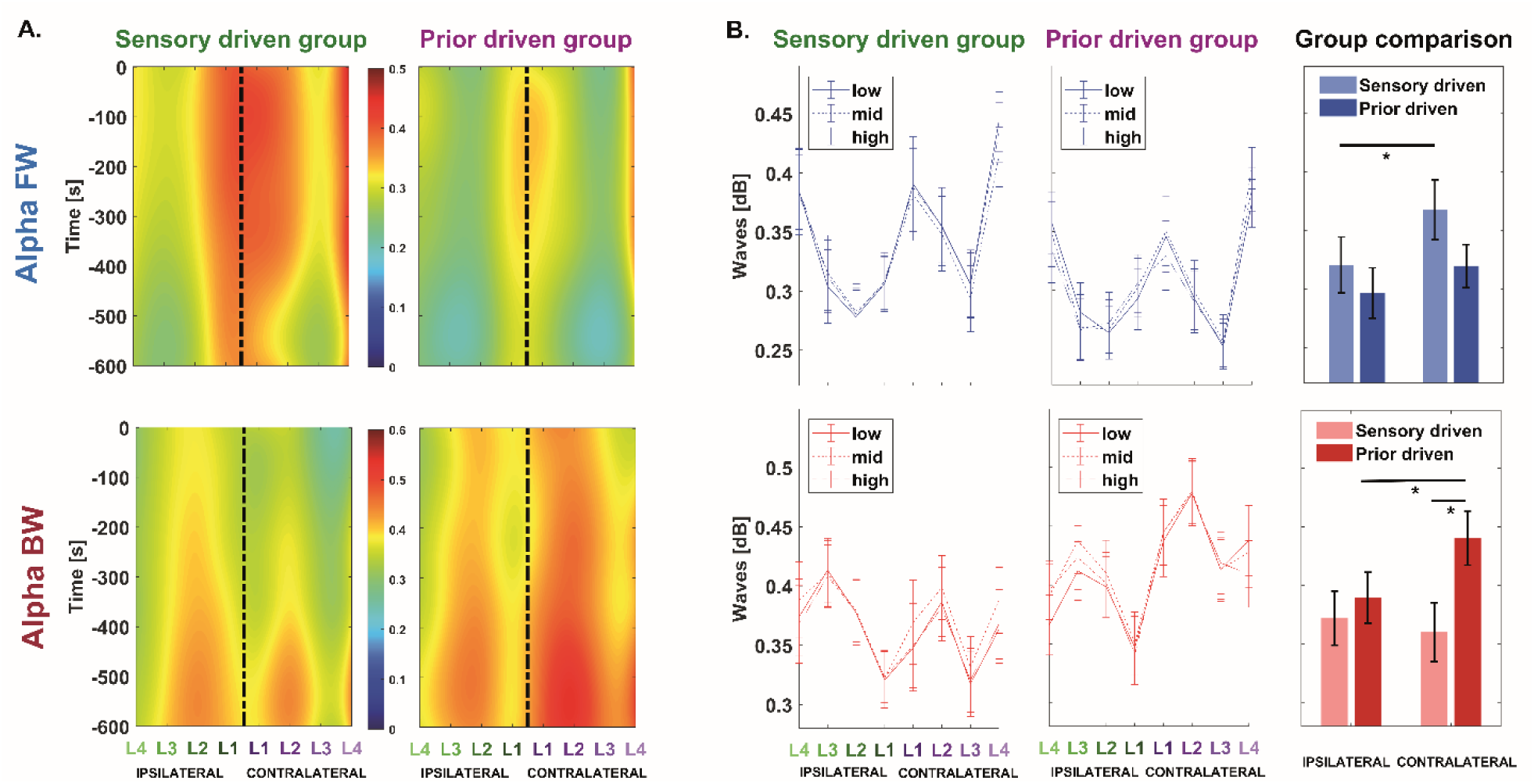
Patterns of travelling waves in prior-driven and sensory-driven individuals. A) We computed forward and backward alpha waves for individuals characterized by prior-driven versus sensory-driven decision-making. The accompanying map depicts the magnitude of these waves during the prestimulus period across the specified electrode lines, with bluish colors indicating low strength and reddish colors representing high strength. B) We observed a hemispheric difference in the magnitude of traveling waves that was moderated by the group factor. Specifically, sensory-driven individuals exhibited a higher prevalence of forward waves in the hemisphere contralateral to stimulus presentation compared to the ipsilateral one. In contrast, prior-driven individuals demonstrated stronger backward waves in the contralateral hemisphere, along with a higher overall magnitude of BW waves compared to the sensory-driven group, particularly in the contralateral hemisphere. This spatial specificity is noteworthy, as the integration of expectations predominantly occurs during the pre-stimulus period in the hemisphere responsible for processing the stimulus. In this study, the right hemisphere played this role, given that the stimulus was consistently presented on the left.

The ANOVA, with BW alpha waves as the dependent variable and within-subject factors HEMISPHERE (contralateral, ipsilateral), LINE (distance from the midline), and CUE (low, neutral, high probability), along with the between-subject factor GROUP (Prior-driven, Sensory-driven), revealed a significant interaction between HEMISPHERE, CUE, and GROUP (F 2,156 = 3.54, p = 0.03, ηp2 = 0.04). Splitting the ANOVA by the GROUP factor revealed no significant effects in the sensory-driven group (all F2,78 < 1.89, all p > 0.16, all ηp2 = 0.05). In contrast, the prior-driven group showed a significant main effect of HEMISPHERE (F = 11.52, p < 0.01, ηp2 = 0.22), indicating that alpha BW waves were more prevalent in the hemisphere contralateral to stimulus presentation (mean alpha BW contralateral= 0.44 ± 0.02 dB) compared to the ipsilateral hemisphere (mean alpha BW ipsilateral= 0.39 ± 0.02 dB). The direct contrast of BW waves between the sensory-driven and prior-driven groups showed that, while both groups exhibited similar levels of BW alpha waves in the ipsilateral hemisphere across all cue levels (all t > -0.66, all p > 0.76, all BF > 0.28), a significant difference emerged in the right hemisphere, with the prior-driven group showing a higher prevalence of BW waves (all t < -1.99, all p < 0.05, BF low probability = 3.24, BF mid probability = 1.28, BF high probability = 2.87). These post hoc results confirm our initial hypothesis, as the observed interaction and subsequent contrasts indicate that the effects of cue probability and group are modulated by hemisphere. Specifically, the contralateral hemisphere shows distinct patterns of BW alpha activity depending on whether the participant adopted a prior-driven or a sensory-driven approach.

Our second hypothesis was partially confirmed, as we observed a trend-level interaction between HEMISPHERE, CUE, and GROUP when analyzing alpha FW waves (F2,156 = 2.63, p = 0.07, ηp2 = 0.03). Although not statistically significant, this finding suggests that the effects of cue probability and hemisphere are modulated by the considered group, in line with our initial predictions. Indeed, further investigation of this interaction, split by the GROUP factor, revealed a pattern opposite to that observed in the BW waves analysis. Specifically, in the prior-driven group, no relationship between cue and hemisphere was found (F2,78 = 0.80, p = 0.45, ηp2 = 0.02). In contrast, a main effect of HEMISPHERE emerged in the sensory-driven group (F1,39 = 8,29, p < 0.01, ηp2 = 0.17), indicating that FW waves were more prominent in the hemisphere contralateral to stimulus presentation (mean FW contralateral = 0.37 ± 0.03) compared to the ipsilateral hemisphere (mean FW ipsilateral = 0.32 ± 0.02) in this group.

Finally, we investigated whether the two groups showed differences in theta and beta oscillations as a function of the other factors tested (see SI). For theta oscillations, the group factor did not show any significant differences, nor did it interact with the other factors, whether considering theta FW or BW waves (all F < 2.68, all p > 0.07, all ηp2 < 0.03). Crucially, a significant interaction between GROUP, LINE, and HEMISPHERE emerged when considering beta BW waves. Follow-up ANOVA, splitting by the GROUP factor, revealed a significant LINE * HEMISPHERE interaction for both the prior- and sensory-driven groups (all F = 3.17, all p < 0.01, all ηp² > 0.09). To further investigate these significant results, we conducted independent t-tests to examine whether the two groups showed significant differences across the four lines as a function of the considered hemisphere. The analysis revealed a significant increase in the BW waves in the left hemisphere, ipsilateral to the presented stimulus hemifield, for the sensory-driven group across all the considered lines (all t39 > 2.21, all p < 0.03, all BF > 1.50), except for the line farthest from the midline (t39 = 1.66, p = 0.10, BF = 0.60). Conversely, the prior-driven group showed an opposite pattern, with stronger beta BW waves in the right hemisphere, contralateral to the presented stimulus hemifield, compared to the left in the two lines closer to the midline (all t39 > 2.69, all p < 0.01, all BF > 3.89), while no differences were observed in the more distant lines (all t39 < 1.48, all p > 0.03, all BF < 0.47).These effects closely resemble, albeit to a lesser strength, those observed within the alpha band.

### Backward waves correlate with alpha-band power modulation

In our previous publication, we demonstrated that the modulation of parieto-occipital alpha amplitude underpins the integration of probabilistic cues into perceptual processing (Figure 4A). Furthermore, Kloosterman et al. ^21^ showed that modulation of posterior alpha amplitude strategically biases evidence accumulation during perceptual tasks. Building on this, we investigated whether the prior-driven effects observed in backward alpha waves were linked to alpha amplitude modulation. We found that contralateral alpha backward traveling waves correlated with the modulation of parieto-occipital alpha power (Figure 4B; Pearson = 0.26, p = ; Spearman = 0.28, p = 0.01; skipped Pearson = 0.24, CI = [0.06, 0.41]; skipped Spearman = 0.27, CI = [0.08, 0.46], BF = 2.06). These results suggest that the magnitude of alpha traveling waves is associated with enhanced modulation of alpha amplitude. Importantly, this relationship was not significant when considering alpha FW waves or BW waves extracted from the ipsilateral hemisphere (Figure 4B; all p > 0.37). Moreover, we demonstrated a positive trial-by-trial relationship between the magnitude of contralateral backward traveling waves and alpha amplitude (See SI), replicating the findings of Alamia et al. ^11^ that revealed the association between these two measures. To further investigate a potential connection between traveling waves, alpha amplitude modulation, and decision criterion adjustments driven by prior cues, we established a mediation model to examine whether the relationship between backward alpha waves and criterion modulation was mediated by concurrent changes in alpha oscillation amplitude (Figure 4C). Crucially, we found a significant mediation effect (Indirect effect = 0.25, CI = [0.06; 0.67]), where an increase in the strength of backward alpha waves is associated with the modulation of decision criterion through a concurrent modulation of alpha amplitude. This suggests that the strength of backward alpha waves plays a key role in shaping decision-making processes by influencing the local alpha power, which in turn affects how prior information is integrated into perceptual judgments.

**Figure 4.**
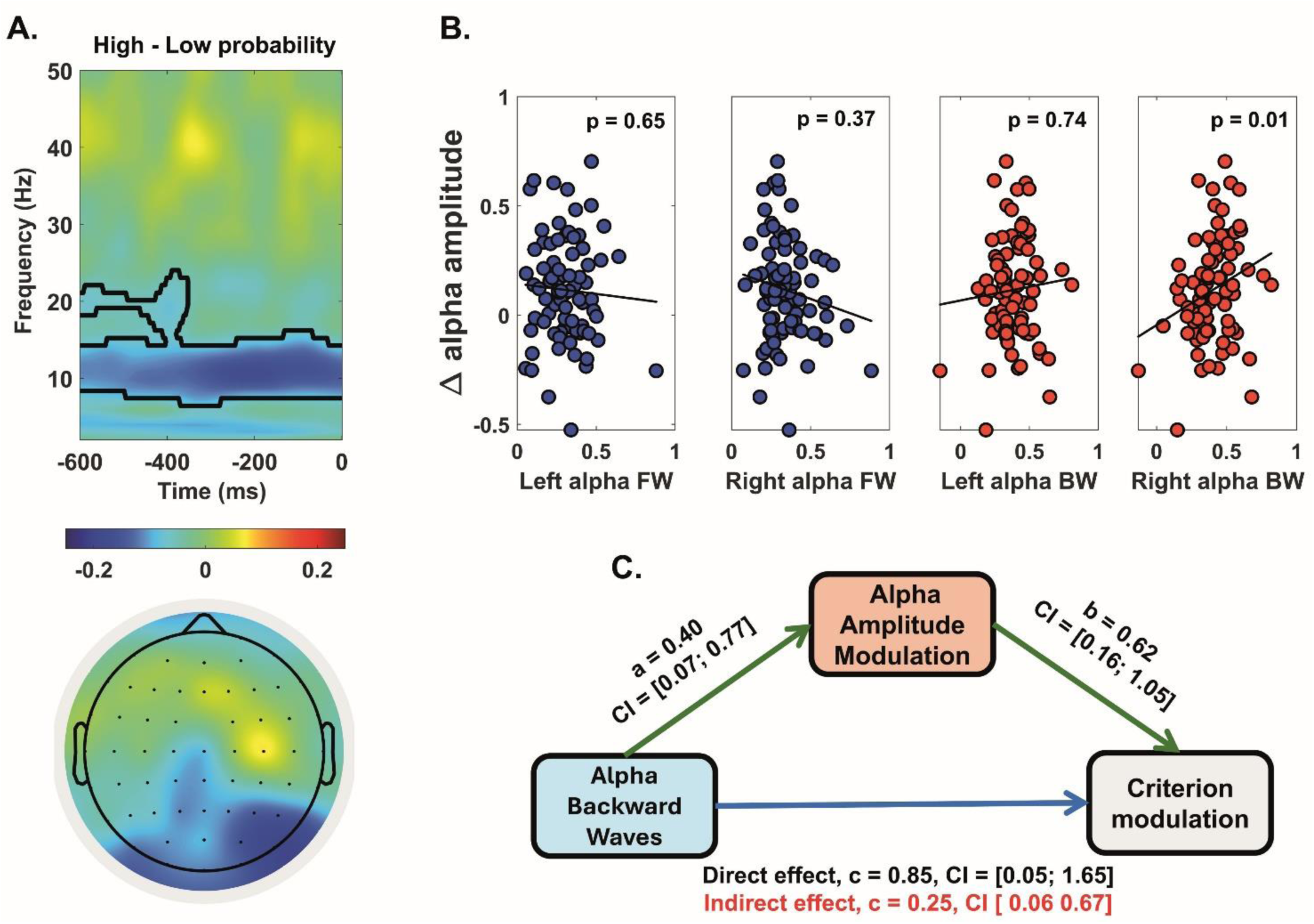
Backward alpha waves modulate the amplitude of occipito-parietal alpha oscillations. A) The images presented here are derived from Tarasi et al., 2022. The top panel shows the time-frequency map of the pre-stimulus period (-600 to 0 ms), highlighting the amplitude differences between high and low probability trials in regions associated with visual processing. Time 0 marks the onset of the stimulus, and black contours indicate statistically significant clusters. The bottom panel depicts the topographic distribution of alpha-band activity differences between low and high probability conditions during the pre-stimulus window. The alpha oscillatory activity diverges between conditions specifically in the posterior electrodes, with a pronounced peak in the right hemisphere. In contrast, the rest of the brain shows comparable activation levels. This highlights a spatially localized effect that modulates cortical activity in areas specialized for stimulus detection. B) While the FW alpha waves and BW waves recorded in the ipsilateral hemisphere did not have a significant effect on the modulation of alpha oscillations, the BW alpha waves recorded in the right hemisphere significantly influenced the alpha amplitude. Specifically, the higher the BW alpha waves, the greater the participant’s ability to modulate the alpha amplitude in the high vs. low probability conditions. C) The mediation analysis demonstrates that traveling waves, alpha amplitude, and decision bias modulation are interrelated. As shown in Tarasi et al., 2022, the degree of modulation of alpha amplitude correlates with the magnitude of decision bias. Similarly, the analyses conducted in this work revealed a significant relationship between the amount of BW waves and the extent of induced bias. Crucially, we show here that the effect linking BW waves and decision bias is mediated by a concurrent modulation of alpha oscillation amplitude. In other words, higher magnitude BW waves are associated with a greater proclivity to modulate alpha amplitude, which, in turn, leads to a more pronounced decision bias.

## DISCUSSION

Previous studies have shown that presenting participants with cues about the likelihood of specific outcomes can significantly bias their decision-making processes. For example, when participants are informed that a target is more likely or unlikely to appear, their rate of reporting the target’s presence increases or decreases accordingly ^5,10,20,31^. These findings demonstrate that cue-based expectations significantly impact perceptual judgments, underscoring the crucial role of prior information in shaping decision outcomes.

In this context, we extend this body of research by highlighting the crucial role of prestimulus alpha traveling waves in integrating prior information during perceptual decision-making processes. Our findings reveal a significant increase in prestimulus backward alpha waves in the contralateral hemisphere (i.e., right), corresponding to the spatial location of the stimulus (i.e., left). This suggests that these waves effectively transmit predictive information to the visual regions responsible for processing the upcoming stimulus. This result supports the predictive coding framework, indicating that the brain uses top-down signals to prepare for anticipated sensory inputs. Furthermore, our findings align with computational models that conceptualize alpha rhythm as the computational code underlying human predictive coding ^33^.

This relationship is further supported by the robust correlation observed between backward alpha waves and prior-driven behavioral modulation in the task: participants exhibiting stronger backward-traveling alpha waves were more likely to adjust their decision criteria according to the probabilistic cues. This highlights the dynamic nature of the alpha rhythm, actively contributing to the integration of expectation-like information into perceptual judgments. In contrast, our results showed no significant correlation between forward alpha waves and prior integration. The absence of an association underscores a functional distinction between the roles of directional alpha waves. Specifically, while backward-traveling waves seem essential for integrating predictive information, forward-traveling waves may serve different functions—likely related to the transmission of processed information through cortical hierarchies, rather than the integration of prior expectations. This result aligns with Alamia et al. ^11^, who demonstrated two functionally distinct traveling waves within the alpha band, propagating in opposite directions during an attentional task: backward waves, which dominate during top-down processes related to attention allocation, and forward waves, which are associated with real-time visual processing during stimulus presentation.

Additionally, we observed a significant correlation between backward waves and modulation of alpha-band power in the occipito-parietal region, consistent with the findings of Kasten et al. ^34^ and Alamia et al. ^11^. Our results support the hypothesis that top-down processes, as reflected by backward waves, drive the established relationship between alpha power and prior information integration, as documented in the literature ^20,21,35–38^. Crucially, mediation analysis corroborated this hypothesis, showing that the increase in the strength of backward alpha waves modulates the decision criterion through a concurrent modulation of alpha amplitude. This suggests a hierarchical process in which the antero-posterior travelling waves, likely reflecting long-range communication of predictive information, modulate the local marker—posterior alpha power—that underlies the integration of prior information. Ultimately, this hierarchically organized mechanism shapes behavioral outcomes that are sensitive to prior cues, as reflected in the modulation of the decision criterion during the task.

Moreover, the magnitude of traveling waves within the alpha band accounted for the differences in predictive styles—specifically, prior-driven individuals (*believers*) versus sensory-driven individuals (*empiricists*)—highlighted in our previous publication within the tested sample (Tarasi et al., 2022). Our findings suggest that only prior-driven participants—those who heavily integrate the cue into the decision-making process—exhibited an increase in backward-traveling waves. In contrast, sensory-driven participants, who rely more on sensory processing for their choices, showed a reduced presence of backward waves and an increased prevalence of forward waves. This weaker anterior-posterior flow suggests that sensory-driven individuals may minimally integrate the prior, as the functional pathway for this integration is under-exploited. In contrast, we observed an increased prevalence of forward-traveling waves in this group, which likely reflects a stronger reliance on sensory processing, where the brain prioritizes the transmission of processed sensory information rather than incorporating predictive cues. In line with previous findings, this pattern would not indicate a fundamental deficit but rather a distinctive integrative style, potentially linked to an autistic-like decision-making approach ^31^, which favors direct stimulus analysis over integrating prior information.

One interesting aspect of our findings is the lateralization of the effects. Specifically, only traveling waves flowing within the right hemisphere exhibited an increase compared to forward waves. Similarly, the increase in backward waves observed in prior-driven versus sensory-driven individuals was specifically found in the right hemisphere. This pattern likely emerged due to the consistent presentation of stimuli in the left visual field. Consequently, the increase in backward alpha activity in the right hemisphere suggests that the brain fine-tunes prior integration by selectively modulating cortical areas involved in processing the target stimuli. This indicates a process carried out with high spatial specificity.

We also investigated whether the observed effect was specific to alpha oscillations or extended to other frequency bands. No significant differences were observed between the prior-driven and sensory-driven groups for FW and BW theta or FW beta waves. However, a difference emerged when examining BW beta waves: the prior-driven group exhibited increased beta activity in the contralateral hemisphere. This finding aligns with evidence suggesting that beta oscillations work in coordination with the alpha band to support the top-down transmission of predictive information _3,4,39._

This work can be expanded through various follow-up studies. For instance, individual differences in the magnitude of these waves could be further investigated, particularly in the context of psychopathological conditions that may exacerbate these mechanisms. Notably, the strong emphasis on priors and idiosyncratic beliefs observed within the schizophrenic spectrum ^40–42^ may be linked to an increased transmission of backward alpha waves ^43,44^. This connection is particularly compelling in light of a recent study ^45^, which found that schizophrenia patients exhibited a substantial increase in top-down alpha-band travelling waves and a decrease in bottom-up waves compared to healthy participants at rest. Investigating this population in tasks similar to the one employed in our study—where high-level explicit priors are provided—could serve as a benchmark for exploring the oscillatory underpinnings of predictive coding disruptions in schizophrenia. An excessive reliance on backward alpha waves might indicate an over-integration of priors, potentially marking a distinct neural signature of this condition.

Moreover, an intriguing avenue for future research could involve establishing an information-based protocol designed to strengthen communication within the anterior-posterior networks through Cortico-Cortical Paired-Associative Stimulation (ccPAS) ^46–50^. ccPAS is a NIBS protocol that leverages Hebbian principles to enhance communication efficiency between networks ^51^. This is achieved by repeatedly pairing TMS pulses delivered to specific nodes within the network at precise, network-dependent timings. By tailoring the timing between the two TMS pulses, it would be possible to achieve modulation in cortical connectivity specific to the frequency of interest (e.g., 100 ms to strengthen alpha wave propagation) ^48,50^. Targeting the anterior-posterior networks involved in conveying predictive activity with this specified timing may facilitate the causal modulation of alpha travelling waves along the predictive chain, thereby enabling an investigation into their causal role in predictive coding.

In conclusion, our study underscores the crucial and frequency-specific role of backward travelling alpha waves in the integration of prior information during perceptual decision-making. By delineating the mechanisms through which expectation-like signals influence perceptual processes, we have contributed to understanding the underlying codes upon which the brain’s predictive machinery relies on.

## Methods

### Participants

Eighty participants (43 female; age range 18-35) provided written informed consent before participating in the study. The study was conducted in accordance with the Declaration of Helsinki and received approval from the Bioethics Committee of the University of Bologna (protocol code 201723, approved on August 26, 2021). The sample is drawn from a previously published dataset _31._

### Stimuli and experimental design

Stimuli were presented on an 18-inch CRT display (resolution 1280 x 1024 pixels, refresh rate 85 Hz) positioned 57 cm away in a dimly lit room. Participants sat comfortably in front of the monitor. The stimuli were generated and displayed using Matlab (version 2016, The MathWorks Inc., Natick, MA) and the Psychophysics Toolbox. The visual stimuli consisted of checkerboards appearing in the lower left visual field, which could either contain grey circles within the cells (target) or not (catch trials). Participants were instructed to indicate the presence (by pressing key ‘k’ with the middle finger) or absence (by pressing key ‘m’ with the index finger) of the grey circles inside the checkerboard as quickly and accurately as possible. To avoid confounding effects related to motor programming, participants responded with their right hand, as this would engage the hemisphere opposite (i.e., the left) to the one responsible for sensory processing in the task (i.e., the right). The study was divided into two phases. In the first, each participant underwent an adaptive titration procedure to determine the contrast of the grey circles for which the detection accuracy was at ∼ 70% when an equal number of target-present and target-absent trials (catch trials) was presented (For detailed explanation of the titration procedure see (Tarasi and Romei, 2024). The second phase comprised 6 blocks of 90 trials each. Each trial started with the appearance of the probability cue presented at the center of the screen. The cue was presented for 1 s followed by a fixation dot. After a variable delay of 1.2–1.5 s a checkerboard containing (or not) grey circles at the titrated contrast within it appeared at the bottom left of the monitor for 60 ms. We opted to present the stimulus in only one hemifields to prevent spontaneous fluctuations in attention between the two hemifields in the prestimulus period from interfering with the results.

Participants had to determine the presence or absence of the grey circles within the checkerboard and press the button associated with their choice. No timeout has been set for the response. After collecting the response, the screen appears black for 1.9-2.4 s in the inter-trial interval. The cue was a rectangle with its bottom colored in red and its top colored in blue. The percentage of the red shading to the entire rectangle indicated the probability that the checkerboard contained the grey circles (target) within it. There were three levels of cues. Cue high and cue low (informative cues) indicated the probability of the presence of the target of 67 and 33%, respectively. Instead, the neutral cue (un-informative cue) equally predicted (50%) the presence and absence of the target. The actual probability of target presentation was in accordance with the probability indicated by the cue. Participants were also explicitly told that the probabilistic cue was congruent with the actual probability of stimulus presentation.

### Signal-detection theory (SDT) modeling

To assess participants’ performance, we applied Signal-Detection Theory (SDT) to compute two key measures: d’ and c ^52^. *d’* represents stimulus sensitivity, with higher values indicating greater sensitivity to the stimulus. *c* represents the decision criterion, where values differing from 0 suggest a bias in decision-making. To determine the influence of probabilistic cues on sensitivity and decision criteria, we computed d’ and c separately for trials that followed low, high, and neutral probability cues. We conducted a repeated-measures ANOVA to examine the effect of cue type on sensitivity and decision criteria, with cue type (high, low, neutral) as the within-subjects factor. To further interpret the results from the ANOVA, we performed post-hoc analyses using paired sample t-tests. Subsequently, for each individual, we calculated the difference in the criterion adopted between conservative and liberal trials (Δ criterion = criterion _low probability trials_ – criterion _high probability trials_) as a measure of cue integration. A larger shift in Δ criterion indicated a greater perceptual adjustment in response to the predictive cue. Conversely, a Δ criterion close to zero suggests minimal perceptual adjustment, indicating that the participant’s decision-making criteria remained stable regardless of the cue provided. All statistical analysis was conducted using the JASP software ^53^.

### EEG analysis

Participants sat comfortably in a dimly lit room. EEG data were collected using a 64-electrode cap, following the international 10–10 system, with signals sampled at 1000 Hz and impedances kept below 10 kΩ. EEG processing was performed offline using custom MATLAB scripts (version R2021a) and the EEGLAB toolbox ^54^. The EEG recordings were filtered in the 0.5-100 Hz range, and a 50 Hz notch filter was applied. The EEG signals were visually inspected, and any noisy channels were spherically interpolated. Epochs from −4100 to 2000 ms relative to the checkerboard onset were extracted. Trials containing excessive noise, muscle, or ocular artifacts were discarded. The recordings were then re-referenced to the average of all electrodes, and Independent Component Analysis (ICA) was applied to remove artifacts distinguishable from brain-driven EEG signals. After artifact removal, the signals were downsampled to 256 Hz

### Traveling wave analysis

We applied a method comparable to that of Alamia et al. ^11^ and Pang et al. ^30^ to assess the propagation of traveling waves across eight lines of seven electrodes, extending from the occipital to frontal regions. As depicted in Figure 1B, we analysed four lines distributed across the left (ipsilateral) and right (contralateral) hemispheres, symmetrically aligned with the midline. The electrode selection overlapped to cover a substantial portion of each hemisphere. For each group of seven electrodes, we generated 2D maps by sliding a 500-ms time window across the EEG signals, with a 125-ms overlap, and computed the 2D-FFT for each map. Importantly, the power in the lower and upper quadrants indicated the extent of waves moving forward (FW – from occipital to frontal electrodes) and backward (BW – from frontal to occipital), respectively. We then repeated the process with shuffled electrode positions to generate a baseline, preserving the spectral power but eliminating directional information (FWss and BWss). Finally, for each frequency between 2 and 45 Hz, we identified the maximum values in the 2D-FFT spectra for both the actual (FW and BW) and shuffled data (FWss and BWss), yielding the wave amount in decibels [dB]. We then calculated the average forward and backward alpha-band waves separately for each line of electrodes and cue.

Finally, we conducted two ANCOVAs, one for each wave direction (forward and backward waves), to evaluate the role of traveling waves in integrating prior information. The analyses included the factors HEMISPHERE (ipsilateral; contralateral), LINE (1 to 4, representing the distance from the midline), and CUE (low, neutral, and high probability cue), with Δ CRITERION as a covariate. The inclusion of the HEMISPHERE factor was essential, as we hypothesized that the effect of traveling waves in prior integration would be specific to the contralateral hemisphere relative to the stimulus presentation. Additionally, the LINE factor was incorporated to explore whether, within each hemisphere, the position along the medial-lateral axis influenced the observed effects. In these ANCOVAs, we focused exclusively on significant effects involving the covariate Δ CRITERION. This choice was driven by the primary aim of the study: to determine how traveling waves contribute to the utilization of perceptual priors and how this process is modulated by cue type, hemisphere, and electrode line position. Any relationships that did not interact with Δ CRITERION were deemed outside the scope of this investigation. We also performed the same analyses on waves extracted from the theta band (5–8 Hz) and beta band (15–25 Hz) to assess the frequency specificity of the effect mediated by the waves in the use of prior information. All ANCOVA results are presented in the supplementary tables.

### Traveling waves and interindividual differences

In our previous publication, we demonstrated that alpha amplitude is a sensitive marker for interindividual differences in the weighting of expectations ^20^. Specifically, we found that a strong modulation of parieto-occipital alpha amplitude between high and low probability conditions was associated with individuals who adopted a prior-driven decision-making strategy. In contrast, individuals who employed a stimulus-oriented approach exhibited no such modulation. Specifically, individuals exhibiting higher alpha power modulation were associated with a prior-driven profile, characterized by a greater criterion shift, while those with lower alpha power modulation were linked to a sensory evidence-driven approach, marked by a reduced criterion shift (see ^20,31^ for further details). Based on the group differentiation identified in previous studies, we conducted two additional ANOVA (one for each direction, FW and BW), with HEMISPHERE (ipsilateral or contralateral), LINE (1 to 4, representing the distance from the midline), CUE (low, neutral, and high probability cue), and GROUP (prior-driven vs. sensory-driven) as factors. Similarly to the approach used in the analyses involving criterion shift, we focused exclusively on effects that demonstrated a significant relationship with the GROUP factor or a significant interaction involving this factor. All other effects were deemed irrelevant to the objectives of the present study, as they did not directly contribute to understanding the role of the adopted predictive strategy in the processing of perceptual priors. To evaluate the frequency specificity of the effects mediated by the waves on the use of prior information, we performed the same analyses on waves extracted from the theta band and beta band. All ANOVA results are presented in the supplementary tables.

### Backward waves and Alpha power modulation

We investigated the relationship between prior-driven effects observed in backward alpha waves and alpha amplitude modulation by conducting Pearson and Spearman correlation analyses, along with their respective skipped versions ^55^, between backward alpha waves and alpha amplitude modulation. We also ensured that the same relationship did not hold when considering FW alpha waves or BW alpha waves recorded in the left hemisphere. Finally, we performed a mediation analysis to explore whether backward alpha traveling waves influenced criterion shifts, with any effects on alpha amplitude modulation serving as a mediator. We report 95% confidence intervals (CI) based on 1000 bootstrap iterations (bias-corrected). Statistical significance was tested by assessing whether 95% of the values excluded zero, as is typically done in this type of analysis ^56^.

Furthermore, we investigated whether there was a significant relationship between alpha traveling waves and alpha amplitude at the single-trial level. This analysis was motivated by previous findings suggesting a crucial link between these two indices (Alamia et al., 2023). To this end, we extracted the magnitude of backward alpha waves at the single-trial level for each electrode line of the contralateral hemisphere. We then computed the average magnitude by collapsing across the electrode lines. Next, we extracted the alpha amplitude again at the single-trial level. For each participant, we correlated the alpha amplitude with the magnitude of the BW waves. Finally, we performed a one-sample t-test against zero on the correlation coefficients to assess whether a significant group-level association existed between the two measures.

## Notes

### Competing Interest Statement

The authors have declared no competing interest.

## REFERENCES

1. Friston, K. & Kiebel, S. Predictive coding under the free-energy principle. Philos Trans R Soc Lond B Biol Sci 364, 1211–1221 (2009).

2. Clark, A. Whatever next? Predictive brains, situated agents, and the future of cognitive science. Behav Brain Sci 36, 181–204 (2013).

3. Bastos, A. M., Lundqvist, M., Waite, A. S., Kopell, N. & Miller, E. K. Layer and rhythm specificity for predictive routing. Proceedings of the National Academy of Sciences 117, 31459–31469 (2020).

4. Bastos, A. M. et al. Visual areas exert feedforward and feedback influences through distinct frequency channels. Neuron 85, 390–401 (2015).

5. Bang, J. W. & Rahnev, D. Stimulus expectation alters decision criterion but not sensory signal in perceptual decision making. Sci Rep 7, 17072 (2017).

6. Lange, F. P. de, Rahnev, D. A., Donner, T. H. & Lau, H. Prestimulus Oscillatory Activity over Motor Cortex Reflects Perceptual Expectations. J. Neurosci. 33, 1400–1410 (2013).

7. Mulder, M. J., Wagenmakers, E.-J., Ratcliff, R., Boekel, W. & Forstmann, B. U. Bias in the brain: a diffusion model analysis of prior probability and potential payoff. J Neurosci 32, 2335–2343 (2012).

8. Tarasi, L., Covelli, M., Fatis, C. T. de & Romei, V. Prior Information Shapes Perceptual Confidence. Journal of Cognition 8, (2025).

9. Tarasi, L. et al. Preparing to act follows Bayesian inference rules. 2024.08.16.608232 Preprint at 10.1101/2024.08.16.608232 (2024).

10. Wyart, V., Nobre, A. C. & Summerfield, C. Dissociable prior influences of signal probability and relevance on visual contrast sensitivity. Proceedings of the National Academy of Sciences 109, 3593–3598 (2012).

11. Alamia, A., Terral, L., D’ambra, M. R. & VanRullen, R. Distinct roles of forward and backward alpha-band waves in spatial visual attention. eLife 12, e85035 (2023).

12. Borghini, G. et al. Alpha Oscillations Are Causally Linked to Inhibitory Abilities in Ageing. The Journal of Neuroscience 38, 4418 (2018).

13. Lobier, M., Palva, J. M. & Palva, S. High-alpha band synchronization across frontal, parietal and visual cortex mediates behavioral and neuronal effects of visuospatial attention. NeuroImage 165, 222–237 (2018).

14. Rihs, T. A., Michel, C. M. & Thut, G. Mechanisms of selective inhibition in visual spatial attention are indexed by alpha-band EEG synchronization. Eur J Neurosci 25, 603–610 (2007).

15. Romei, V., Rihs, T., Brodbeck, V. & Thut, G. Resting electroencephalogram alpha-power over posterior sites indexes baseline visual cortex excitability. NeuroReport 19, 203 (2008).

16. Romei, V. et al. Spontaneous Fluctuations in Posterior α-Band EEG Activity Reflect Variability in Excitability of Human Visual Areas. *Cerebral Cortex (New York*, NY) 18, 2010 (2007).

17. Thut, G., Nietzel, A., Brandt, S. A. & Pascual-Leone, A. Alpha-band electroencephalographic activity over occipital cortex indexes visuospatial attention bias and predicts visual target detection. J Neurosci 26, 9494–9502 (2006).

18. Trajkovic, J., Di Gregorio, F., Avenanti, A., Thut, G. & Romei, V. Two Oscillatory Correlates of Attention Control in the Alpha-Band with Distinct Consequences on Perceptual Gain and Metacognition. J Neurosci 43, 3548–3556 (2023).

19. Alamia, A. & VanRullen, R. Alpha oscillations and traveling waves: Signatures of predictive coding? PLOS Biology 17, e3000487 (2019).

20. Tarasi, L., di Pellegrino, G. & Romei, V. Are you an empiricist or a believer? Neural signatures of predictive strategies in humans. Progress in Neurobiology 219, 102367 (2022).

21. Kloosterman, N. A. et al. Humans strategically shift decision bias by flexibly adjusting sensory evidence accumulation. eLife 8, e37321 (2019).

22. Limbach, K. & Corballis, P. M. Prestimulus alpha power influences response criterion in a detection task. Psychophysiology 53, 1154–1164 (2016).

23. Benwell, C. S. Y., Coldea, A., Harvey, M. & Thut, G. Low pre-stimulus EEG alpha power amplifies visual awareness but not visual sensitivity. Eur J Neurosci 55, 3125–3140 (2022).

24. Di Gregorio, F. et al. Tuning alpha rhythms to shape conscious visual perception. Current Biology 32, 988–998.e6 (2022).

25. Trajkovic, J., Di Gregorio, F., Thut, G. & Romei, V. Transcranial magnetic stimulation effects support an oscillatory model of ERP genesis. Current Biology 34, 1048–1058.e4 (2024).

26. Petras, K., Grabot, L. & Dugué, L. Locally induced traveling waves generate globally observable traveling waves. 2025.01.07.630662 Preprint at 10.1101/2025.01.07.630662 (2025).

27. Sato, T. K., Nauhaus, I. & Carandini, M. Traveling Waves in Visual Cortex. Neuron 75, 218–229 (2012).

28. Zhang, H., Watrous, A. J., Patel, A. & Jacobs, J. Theta and Alpha Oscillations Are Traveling Waves in the Human Neocortex. Neuron 98, 1269–1281.e4 (2018).

29. Mohanta, S. et al. Traveling waves shape neural population dynamics enabling predictions and internal model updating. 2024.01.09.574848 Preprint at 10.1101/2024.01.09.574848 (2024).

30. Pang, Z., Alamia, A. & VanRullen, R. Turning the Stimulus On and Off Changes the Direction of α Traveling Waves. eNeuro 7, (2020).

31. Tarasi, L., Martelli, M. E., Bortoletto, M., di Pellegrino, G. & Romei, V. Neural Signatures of Predictive Strategies Track Individuals Along the Autism-Schizophrenia Continuum. Schizophr Bull 49, 1294–1304 (2023).

32. Tarasi, L. & Romei, V. Individual Alpha Frequency Contributes to the Precision of Human Visual Processing. Journal of Cognitive Neuroscience 36, 602–613 (2024).

33. Schwenk, J. C. B. & Alamia, A. A hierarchical multiscale model of forward and backward alpha-band traveling waves in the visual system. 2024.11.15.623743 Preprint at 10.1101/2024.11.15.623743 (2024).

34. Kasten, F. H., Wendeln, T., Stecher, H. I. & Herrmann, C. S. Hemisphere-specific, differential effects of lateralized, occipital–parietal α-versus γ-tACS on endogenous but not exogenous visual-spatial attention. Sci Rep 10, 12270 (2020).

35. Albu, S. & Meagher, M. W. Expectation of nocebo hyperalgesia affects EEG alpha-activity. Int J Psychophysiol 109, 147–152 (2016).

36. Rohenkohl, G. & Nobre, A. C. α oscillations related to anticipatory attention follow temporal expectations. J Neurosci 31, 14076–14084 (2011).

37. Samaha, J., Boutonnet, B., Postle, B. R. & Lupyan, G. Effects of meaningfulness on perception: Alpha-band oscillations carry perceptual expectations and influence early visual responses. Sci Rep 8, 6606 (2018).

38. Sáringer, S., Fehér, Á., Sáry, G. & Kaposvári, P. Perceptual Expectations Are Reflected by Early Alpha Power Reduction. Journal of Cognitive Neuroscience 36, 1282–1296 (2024).

39. Clayton, M. S., Yeung, N. & Cohen Kadosh, R. The many characters of visual alpha oscillations. European Journal of Neuroscience 48, 2498–2508 (2018).

40. Haarsma, J. et al. Influence of prior beliefs on perception in early psychosis: Effects of illness stage and hierarchical level of belief. Journal of Abnormal Psychology 129, 581–598 (2020).

41. Stuke, H., Kress, E., Weilnhammer, V. A., Sterzer, P. & Schmack, K. Overly Strong Priors for Socially Meaningful Visual Signals Are Linked to Psychosis Proneness in Healthy Individuals. Front. Psychol. 12, (2021).

42. Tarasi, L., Borgomaneri, S. & Romei, V. Antivax attitude in the general population along the autism-schizophrenia continuum and the impact of socio-demographic factors. Front. Psychol. 14, (2023).

43. Ippolito, G. et al. The Role of Alpha Oscillations among the Main Neuropsychiatric Disorders in the Adult and Developing Human Brain: Evidence from the Last 10 Years of Research. Biomedicines 10, 3189 (2022).

44. Tarasi, L. et al. Predictive waves in the autism-schizophrenia continuum: A novel biobehavioral model. Neurosci Biobehav Rev 132, 1–22 (2022).

45. Alamia, A. et al. Oscillatory traveling waves provide evidence for predictive coding abnormalities in schizophrenia. Biological Psychiatry (2024) doi:10.1016/j.biopsych.2024.11.014.

46. Luzio, P. D., Tarasi, L., Silvanto, J., Avenanti, A. & Romei, V. Human perceptual and metacognitive decision-making rely on distinct brain networks. PLOS Biology 20, e3001750 (2022).

47. Romei, V., Chiappini, E., Hibbard, P. B. & Avenanti, A. Empowering Reentrant Projections from V5 to V1 Boosts Sensitivity to Motion. Curr Biol 26, 2155–2160 (2016).

48. Romei, V., Thut, G. & Silvanto, J. Information-Based Approaches of Noninvasive Transcranial Brain Stimulation. Trends in Neurosciences 39, 782–795 (2016).

49. Sel, A. et al. Increasing and decreasing interregional brain coupling increases and decreases oscillatory activity in the human brain. Proceedings of the National Academy of Sciences 118, e2100652118 (2021).

50. Tarasi, L., Turrini, S., Sel, A., Avenanti, A. & Romei, V. Cortico-cortical paired-associative stimulation to investigate the plasticity of cortico-cortical visual networks in humans. Current Opinion in Behavioral Sciences 56, 101359 (2024).

51. Caporale, N. & Dan, Y. Spike timing-dependent plasticity: a Hebbian learning rule. Annu Rev Neurosci 31, 25–46 (2008).

52. Green, D. M. & Swets, J. A. *Signal Detection Theory and Psychophysics*. xi, 455 (John Wiley, Oxford, England, 1966).

53. Love, J. et al. JASP: Graphical Statistical Software for Common Statistical Designs. Journal of Statistical Software 88, 1–17 (2019).

54. Delorme, A. & Makeig, S. EEGLAB: an open source toolbox for analysis of single-trial EEG dynamics including independent component analysis. J Neurosci Methods 134, 9–21 (2004).

55. Pernet, C. R., Wilcox, R. & Rousselet, G. A. Robust correlation analyses: false positive and power validation using a new open source matlab toolbox. Front Psychol 3, 606 (2012).

56. Hayes, A. Introduction to Mediation, Moderation, and Conditional Process Analysis: Third Edition: A Regression-Based Approach. Guilford Press https://www.guilford.com/books/Introduction-to-Mediation-Moderation-and-Conditional-Process-Analysis/Andrew-Hayes/9781462549030 (2022).

